# A provably convergent control closure scheme for the Method of Moments of the Chemical Master Equation

**DOI:** 10.1101/2023.05.03.539185

**Authors:** Vincent Wagner, Robin Strässer, Frank Allgöwer, Nicole Radde

## Abstract

In this article, we introduce a novel moment closure scheme based on concepts from Model Predictive Control (MPC) to accurately describe the time evolution of the statistical moments of the solution of the Chemical Master Equation (CME). The Method of Moments, a set of ordinary differential equations frequently used to calculate the first *n*_m_ moments, is generally not closed since lower-order moments depend on higher-order moments. To overcome this limitation, we interpret the moment equations as a nonlinear dynamical system, where the first *n*_m_ moments serve as states and the closing moments serve as control input. We demonstrate the efficacy of our approach using three example systems and show that it outperforms existing closure schemes. For polynomial systems, which encompass all mass-action systems, we provide probability bounds for the error between true and estimated moment trajectories. We achieve this by combining convergence properties of a priori moment estimates from stochastic simulations with guarantees for nonlinear reference tracking MPC. Our proposed method offers an effective solution to accurately predict the time evolution of moments of the CME, which has wide-ranging implications for many fields, including biology, chemistry, and engineering.

## 1 Introduction

The dynamics of well-stirred chemical reaction systems are described by the Chemical Master Equation (CME). The CME is a set of linear differential equations for the time-dependent probability distribution over the set of all possible configurations of the system. Since this set is usually large or even infinite except for very small systems, solving the CME directly is often not feasible. One remedy for this problem is to consider only the first *n*_m_ moments of this probability distribution, thereby reducing the number of equations from the number of configurations to the number of considered moments.

Two established methods to estimate the statistical moments of the CME solution are the Stochastic Simulation Algorithm (SSA) and the Method of Moments (MoM). The SSA (1, 2) generates sample paths from the underlying Markov process, which can be used to approximate either the full distribution or its moments via Monte Carlo integration. SSA-based moment approximation has several advantages. First, it inherently produces physically meaningful moment estimates, for example, means and variances are non-negative. Second, empirical moment estimates are consistent and thus converge in probability to the true moments in the large sample limit. On the other hand, approximating the CME solution or its moments only based on SSA samples is often tedious, as the computational load of generating each sample grows with the number of simulated molecules. Moreover, a huge number of samples is needed for an accurate estimate if, for example, some species have a high probability of becoming zero, if the system is stiff, or if it includes rare events.

The MoM is an infinite set of coupled Ordinary Differential Equations (ODEs) which describes the evolution of the statistical moments of the CME solution. Mathematically, the equation for each moment is obtained by multiplying the CME with a moment-specific test function and summation over all configurations, as described in (3). The solution of this MoM ODE system can be obtained with state-of-the-art solvers and the needed computational power is nearly independent of the number of simulated molecules. However, one challenging aspect of the MoM ODEs is their interdependence structure. For systems with quadratic propensities, a moment of a given order depends on the next higher-order moment, leading to an infinite dependency hierarchy, and the set of equations for the first *n*_m_ moments is not closed. It is worth mentioning that closing the MoM equations for the first *n*_m_ moments with the true higher-order moments, would they be known hypothetically, results in exact moment estimates. Plenty of solution approaches have been proposed to solve this closure problem. We will briefly outline some of them and refer the interested reader to (4) for an in-depth review of the general moment closure problem.

The most trivial closure technique simply neglects the influence of all higher-order moments on the considered lower-order moments and is thus effectively a truncation of the ODE system. We, therefore, refer to it as Truncation Closure (TC). TC is trivially implementable without any additional manual or computational effort. Further closure techniques have been introduced for stochastic differential equation systems (5), including the cumulant closure technique that was first proposed by Cramer (6). Cumulants are, like moments, characteristic quantities of a distribution that can be calculated via its characteristic function. Similar to TC, the cumulant closure technique closes the cumulant equations by setting all higher-order cumulants to zero. Since the first two cumulants correspond to expectation value and variance, respectively, cumulant closure is equivalent to TC if only the dynamics of the first two moments are considered.

Beyond these naïve approaches, a popular class of methods assumes that the CME solution is similar to a specific family of distributions and uses the higher-order moments of these respective distributions for closure (7–12). Other approaches express higher-order moments as potentially nonlinear functions of the considered lower-order moments (13–18). A third interesting class reconstructs the underlying distribution at one or several time steps, e.g., by exploiting the maximum entropy principle, and uses the reconstruction to infer the closure moments (19–21).

Many of the proposed closure schemes work very well in practice. For special cases, there are even theoretical results justifying specific closure schemes (22–24). However, since there is generally no relation between the trajectories of the closed moment equations and the CME, both are decoupled. Hence, moment closure methods suffer from undesired artifacts such as unphysical moments and potentially large deviations from the true moment trajectories, which constitutes the major drawback of closure methods. Even worse, except for systems with only reactions of up to order one, for which the moment equations are naturally closed, we do not have criteria at hand which allow for an a priori judgment of the quality of a moment closure solution based on the set of chemical reactions, the initial species amounts and number *n*_m_ of considered moments. Therefore, a closure scheme that generally and provably closes the MoM ODEs in accordance with the true CME solution would be highly desirable since it solves this major problem.

In this article, our aim is to address this decoupling issue by providing a closure that guarantees that the distance between the moment closure solutions and the true moments is bounded and moments are physically plausible. To achieve this ambitious goal, we employ a few SSA samples that are responsible for coupling the closed MoM ODEs to the true CME solution. This idea is related to the work in (25) that uses a few SSA trajectories to roughly estimate the closure moments and subsequently uses Kalman filtering to eradicate the noise. While their proposed closure method relies on *sampled* closing moments that are typically affected by non-negligible noise, we propose to obtain the closing moments as a control input. To this end, we interpret the closed moment equations as a nonlinear system, in which the moments up to order *n*_m_ serve as states and the closing moments serve as a control input. For this *Control Closure (CC) scheme*, we can build on a variety of controller design methods for nonlinear systems from the literature (26). Particularly for the class of polynomial dynamics, which contains for example all mass-action systems, a popular approach for non-linear control relies on sum-of-squares optimization, which reformulates non-negativity conditions as semi-definite programs (27, 28). However, sum-of-squares optimization is generally computationally challenging. Instead, we rely on Model Predictive Control (MPC) (29, 30) and, in particular, use concepts of nonlinear reference tracking MPC as presented in (31, 32). For CC, we characterize SSA-based estimates and confidence intervals of the moments by exploiting concentration inequalities of random variables. We then combine these with the MoM by formulating an optimal control problem in which the resulting optimal control input serves as the proposed moment closure. We validate our approach on three example systems, for which we obtain very good performance compared to TC and stand-alone SSA estimation. In particular, the higher-order moment estimates obtained by our CC scheme are indeed close to the true moments. While there are other more sophisticated closure techniques, e.g., Gaussian closure (13), as discussed previously, we restrict ourselves to the comparison to TC as the simplest closure method at hand which is also widely used in practice. The particular contribution of the present paper is not limited to the proposed generally applicable CC and its sheer comparison with existing closure techniques but includes a theoretical analysis of the proposed closure. In particular, for polynomial systems, we provide guarantees that the moment trajectories inferred from CC satisfy the moment closure equations and lie in the given confidence intervals while staying close to the sampled moment estimates.

### Notation

Throughout this paper, bottom right indices _*i*_ denote entries of vector-valued quantities. Samples for example from the SSA are indexed using a top left index in parentheses ^(*i*)^ . To sum over different chemical reactions or system configurations, we employ top right indices in parentheses ·^(*i*)^. For a symmetric matrix *A*, we write *A* ≻ 0 (*A* ⪰ 0) if *A* is positive (semi-)definite. We write ∥*x*∥_*A*_ for the weighted norm *x*^⊤^*Ax* of a vector *x* with *A* ≻ 0. By ℕ_0_ and ℝ_*≥*0_ we denote the natural numbers including zero and all nonnegative real numbers, respectively. We use the vectorial index 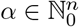 with |*α*| = *α*_1_ + … + *α*_*n*_ to define for a vector *x* ∈ ℝ^*n*^ the monomial 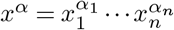. We denote the class of functions *α* : ℝ_*≥*0_ → ℝ_*≥*0_, which are continuous, strictly increasing, and satisfy *α*(0) = 0 by 𝒦. We write 𝒦_*∞*_ for the set of functions *α* ∈ 𝒦 satisfying lim_*k→∞*_ *α*(*k*) = ∞. By ℒ we denote the class of functions *β* : ℝ_*≥*0_ → ℝ_*≥*0_, which are continuous and decreasing with lim_*k→∞*_ *β*(*k*) = 0. A continuous function *β* : ℝ_*≥*0_ × ℝ_*≥*0_ belongs to class 𝒦 ℒ if *β*(·, *s*) ∈ K for each fixed *t* and *β*(*r*, ·) ∈ ℒ for each fixed *r*.

## 2 SSA trajectories as control reference for moment closure

In this section, we present a CC scheme to close the moment equations of the CME. Subsection 2-A first introduces an interpretation of the closed moment equations from the perspective of control theory. Second, we derive probability bounds for SSA-based moment estimation in Subsection 2-B, which define a confidence region that contains the true moments with a certain user-defined probability. Subsequently, we use these SSA-based estimates and probability bounds in Subsection 2-C to formulate an optimization problem whose solution is the best possible estimate of the moments according to a predefined optimization criterion, while satisfying the closed moment equations, staying close to the SSA-based moment estimates, and lying inside the confidence region. Since this optimization problem tends to be computationally demanding for large time intervals, we present a short-horizon tracking MPC scheme for which we develop theoretical results leading to a provably convergent CC for optimal estimation of the true CME moments in Section 2-D.

### A. System theoretic interpretation of the closed moment equations

The CME is a set of linear ordinary differential equations that describes the time-dependent probability *p*(*z, t*) for each system configuration *z* as

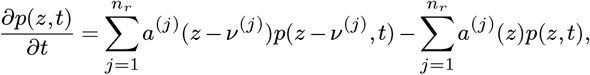

with reaction propensities *a*^(*j*)^(*z*), configuration change vectors *ν*^(*j*)^, and sums over all feasible reactions. The set Ω_*Z*_ of feasible configurations depends on the set of reactions and the initial molecule numbers. For a simple reversible reaction *A* ⇌ *B* and a single molecule, for instance, Ω_*Z*_ = {*z*^(1)^ = (1, 0), *z*^(2)^ = (0, 1)}. Moment equations are obtained by multiplication with a moment-specific test function *T* (*z*) : Ω_*Z*_→ ℝ and summing over all configurations (3), i.e.,

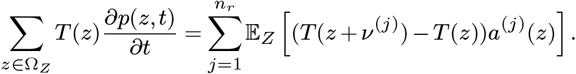

For example, for *T* (*z*) = *z*_*i*_ and *T* (*z*) = (*z*_*k*_ − E_*Z*_(*z*_*k*_))(*z*_*l*_ − E_*Z*_(*z*_*l*_)), respectively, we get for the expectation 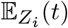 and the covariance 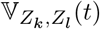 between *Z*_*k*_ and *Z*_*l*_

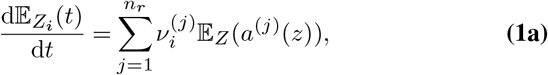

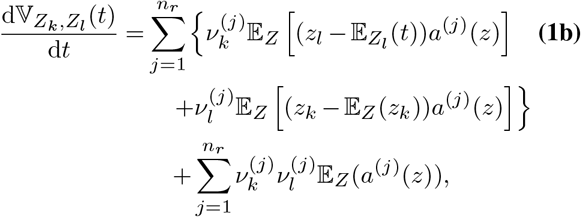

with 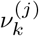 denoting the *k*-th component of the vector *ν*^(*j*)^ (3). The general moment equations Eq. (1) can be interpreted as a controlled system of the form

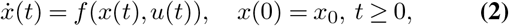

where *x* ∈ R^*n*^ contains the first *n*_m_ moments that appear as states and *u* ∈ ℝ^*m*^ defines the control input and corresponds to the higher-order moments that are needed to close the system. Further, we write *x*_*u*_(*t*; *x*_0_) for the solution of Eq. (2) for a given control sequence *u* and initial condition *x*_0_. The proposed CC scheme is in principle valid for arbitrary dimensions and moment orders, and we later also apply it to a system with two chemical species to show its broad applicability. However, for ease of notation, we restrict the following derivations to systems with only one chemical species (*n*_c_ = 1) and refer the interested reader to Engblom (3) for a general derivation of the MoM. We are interested in predicting the expectation value and variance (*n*_m_ = 2), i.e.,

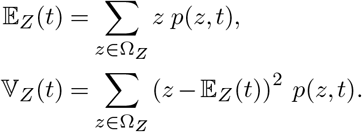

In this case, a second order Taylor series expansion of *a*^(*j*)^(*z*_0_) about *z*_0_ = E_*Z*_(*t*) in Eq. (1) leads to (see Appendix A)

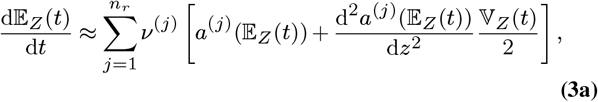

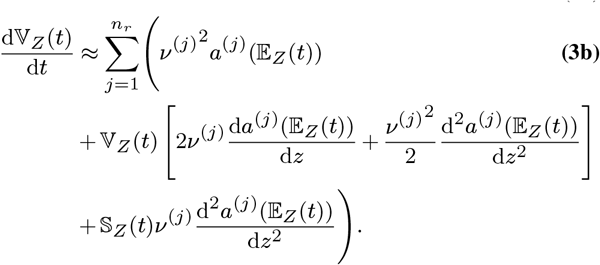

In this representation, the skewness

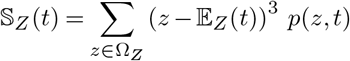

is the closure moment.

For mass-action systems with reactions of at most quadratic propensities, the MoM ODEs are at most quadratic in the moments, i.e., there are no products of more than two moments. Moreover, the moments of order (*n*_m_ + 1) affect only the *n*_m_-th moments and occur only linearly in the corresponding ODE (3). In this case, the Taylor expansion Eq. (3) is exact and can be used for further specification of the dynamical system Eq. (2) to

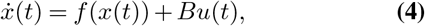

where 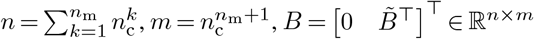 with 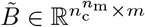, and *f* : ℝ^*n*^ → ℝ^*n*^ is a quadratic drift term, i.e.,

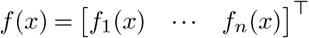

with

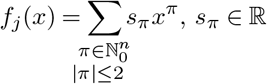

for all *j* = 1, …, *n*. For *n*_c_ = 1 and *n*_m_ = 2, the dynamical systems Eq. (2) and Eq. (4) are of dimension *n* = 2, the closing moment *u* is of dimension *m* = 1, and the matrix *B* in Eq. (4) reduces to a vector, i.e., 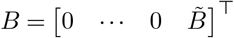 with 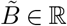. With this simplification, we finally have a two-dimensional controlled system for the expectation value 𝔼_*Z*_(*t*) and the variance 𝕍_*Z*_(*t*), which reads

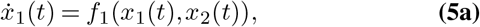

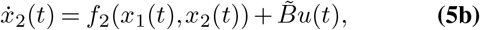

where *f*_1,2_(*x*_1_, *x*_2_) are polynomial functions in their arguments. In the following, we use the derived dynamical system representation to formulate an optimization-based closure for the MoM. To this end, we rely on SSA to obtain rough moment estimates together with probabilistic bounds on their accuracy.

### B. Probability bounds for SSA-based moment estimation

Empirical moment estimates 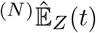 from SSA sample paths ^(*i*)^*z*(*t*), *i* = 1, …, *N*, are consistent and converge for each time instance *t*^*^ in probability to the true moment 𝔼_*Z*_(*t*^*^) for large sample sizes, i.e.,

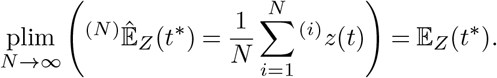

In an attempt to quantify this convergence for a finite number of SSA trajectories *N*, we are interested in a probabilistic bound for 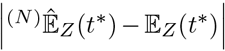, or in other words, confidence intervals of the mean 𝔼_*Z*_(*t*^*^). Estimating these intervals is one of the most fundamental questions of frequentist statistics and consequently, the amount of related literature is vast. Ideally, we would like to present a probabilistic bound of the form

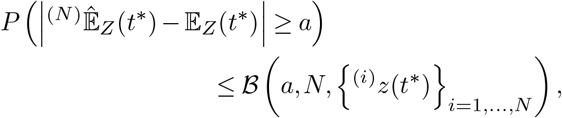

where the bounding function ℬ only depends on the desired width *a >* 0 of the confidence interval, the number of drawn samples *N*, and the samples themselves. However, it seems to be impossible to obtain such a purely sample-based bound ℬ. This subsection, therefore, outlines the most prominent approaches together with their respective additional assumptions and refers the interested reader to relevant sources for more information. We would like to stress that the proposed CC scheme does not rely on any particular bound. Thus, every applicant of our technique is able to choose bounds according to each individual situation.

One of the most prominent representatives of this class of inequalities is the Chebyshev-Inequality (CI)

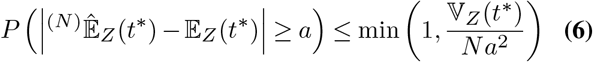

for all *a* ∈ ℝ, or closely related variants thereof (33). However, its bound depends on the unknown *true* variance 𝕍_*Z*_(*t*^*^).

In a similar fashion, the Central Limit Theorem (CLT) can be combined with the Berry-Esseen theorem (34, 35) to show

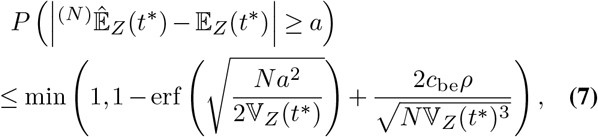

which is detailed in Appendix B. Here, *c*_be_ is the Berry-Esseen constant that is known to be upper bounded by 0.4748 (36), erf denotes the error function, and *ρ* is the *true* third centralized absolute moment

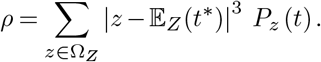

Again, knowledge of the *true* variance 𝕍_*Z*_(*t*^*^) and the just-defined moment *ρ* is necessary to obtain the desired bound. In contrast, Hoeffding’s Inequality (HI) (37) provides a bound independent of higher-order moments but instead requires bounded states 0 ≤ ^(*i*)^*z*(*t*) ≤ *b, b* ∈ ℝ. It reads

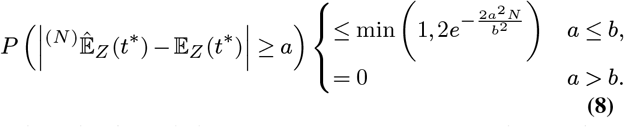

While this boundedness is a strong assumption, the number of system molecules is often bounded by conservation laws anyway. In case the system can indeed grow infinitely, a configuration state truncation can provide such an upper bound *b*, which is, however, not uniquely defined. Both named possibilities will be demonstrated in our example systems.

By defining an upper bound *α* ∈ (0, 1) for the probability in one of the inequalities Eq. (6), Eq. (7), and Eq. (8), each inequality can be resolved for the parameter *a*. In case of the HI, this leads to

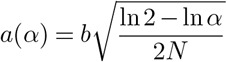

exemplary, and the true moment lies with probability 1 − *α* in the symmetric interval 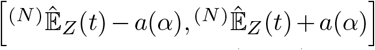 around the SSA-estimate, which serves as (1 − *α*) confidence interval. In addition, physical constraints such as the non-negativity of expectation and variance might be used to tighten these intervals even further if desired. We denote the union of confidence intervals for all *t*^*^ in a given time interval 𝒯:= [0, *T*_end_] with *T*_end_ *>* 0 as confidence region, which describes a symmetric tube around 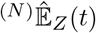 for *t* ∈ 𝒯. For a bounded state space, also the variance could be constrained. While we restricted this subsection to bounds for 𝔼_*Z*_(*t*^*^), many results can be generalized to higher-order moments and similarly included in the CC scheme presented later. For these and many other related results, we can (upon many others) recommend the textbook of (38) and the articles of (33), (39), and (40).

### C. Optimal CC scheme for CME moment estimation

Based on the system theoretic representation Eq. (4) of the moment equations as a dynamical system, we formulate a CC scheme that relies on a priori SSA stand-alone moment estimates and confidence regions derived in Subsection 2-B and uses MPC concepts to find an optimal input *u*. In particular, we formulate an optimization problem to find a control input *u* which guarantees that the MoM with CC leads to estimated moments that minimally deviate from the SSA-based moment estimation 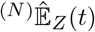 with respect to a specified cost function and are guaranteed to lie within the predefined confidence region.

A graphical overview of the proposed CC is depicted in Figure 1. Given SSA trajectories ^(*i*)^*z*(*t*) over the finite time interval 𝒯, we obtain estimates of the first (*n*_m_ + 1) moments. We collect the first *n*_m_ moments in 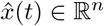 and all moments of order (*n*_m_ + 1) in *û*(*t*) ∈ ℝ^*m*^ for *t* ∈ 𝒯 . Where applicable, we then define the constraint sets 𝒳 (*t*) ⊆ℝ^*n*^ and 𝒰(*t*) ⊆ℝ^*m*^ as the SSA-based confidence regions in Subsection 2-B. We want to ensure (*x*(*t*), *u*(*t*)) ∈ 𝒳 (*t*) × 𝒰(*t*) for all *t*∈ 𝒯. Both 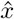 and *û* as well as 𝒳(*t*) and 𝒰(*t*) influence the MPC controller, which is explained in the following and returns moment estimates 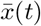 and 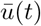 that lie within 𝒳(*t*) and 𝒰(*t*).

**Fig. 1.**
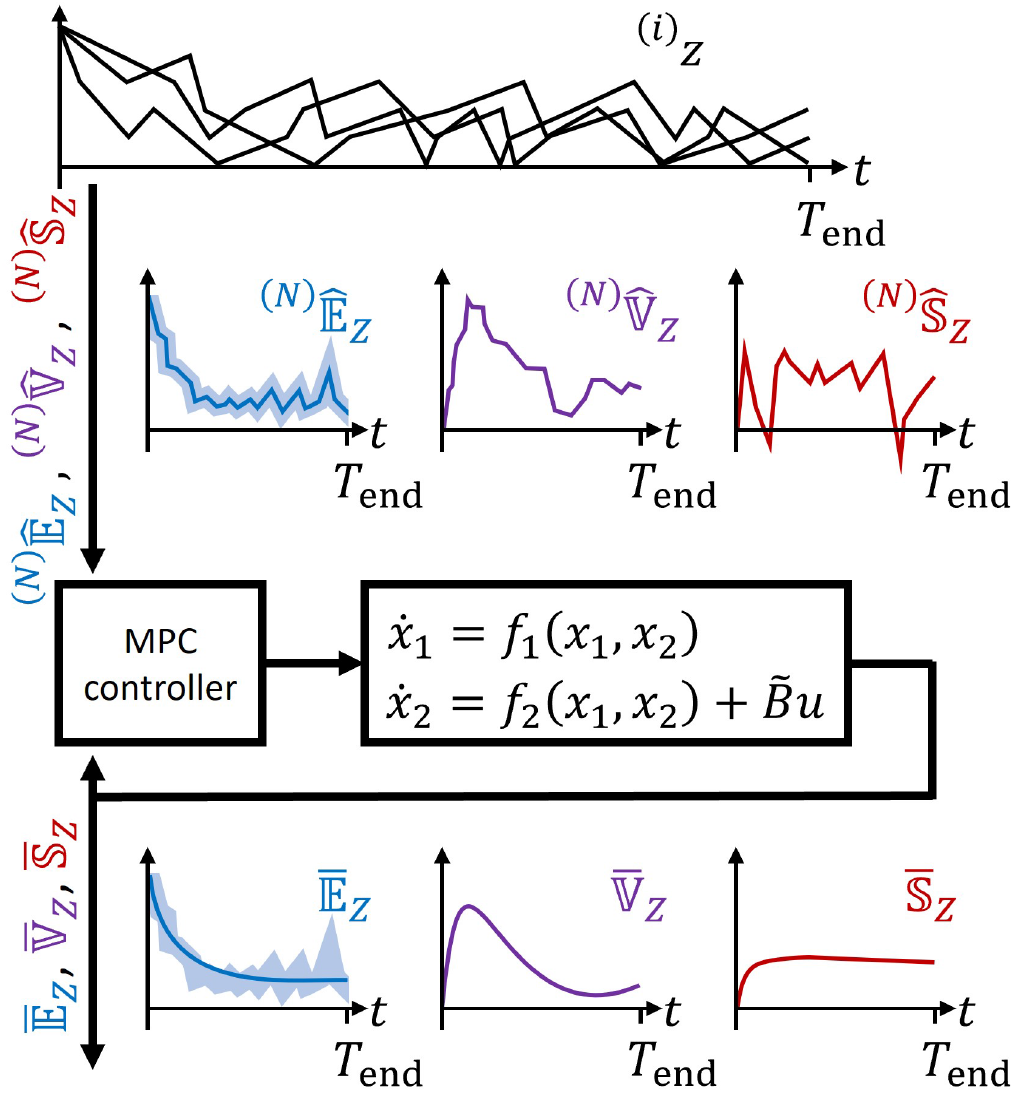
Control closure scheme for one chemical species and two considered moments. According to the dynamics in Eq. (5), the state *x* ∈ ℝ^2^ consists of expectation and variance, and the skewness, which constitutes the input *u*(*t*), appears in *f*_2_ as a linear term with factor 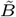. The controller takes SSA-based estimations 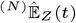 and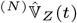 and, if applicable, respective confidence regions as inputs and uses an MPC algorithm that returns moment estimates 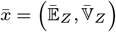, and 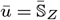. The time indication is omitted for brevity.

As we seek the optimal moment estimation on the interval 𝒯, we need to define an optimality criterion first. To this end, we define the open-loop cost function

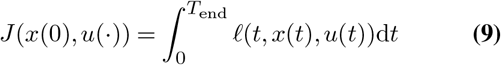

with the stage cost

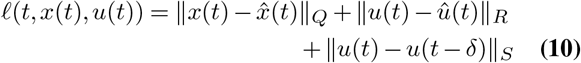

for some user-defined positive definite *Q* ∈ ℝ^*n×n*^, positive semi-definite *R, S* ∈ ℝ^*m×m*^, and sampling period *δ >* 0. These matrices can be chosen, e.g., to weigh some moments more than others. If there is no estimate *û*(*t*) of the closing moments, we can just choose *R* = 0 or *û*(*t*) = 0. The last term in Eq. (10) is used to penalize the rate of change of the resulting input *u*. The matrix *S* can be seen as a tuning parameter: For a larger weight *S*, the difference in *u* at two consecutive time points *t* − *δ* and *t* is penalized more, and, hence, the optimization yields a smoother trajectory for the control input.

#### Definition 1

For weights *Q* ≻ 0, *R* ⪰ 0, and *S* ⪰ 0 and an initial condition *x*(0) = *x*_0_, the optimal reachable trajectory (*x*^*^(*t*), *u*^*^(*t*)) on the interval 𝒯 is the minimizer of

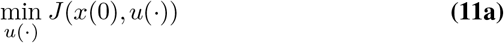

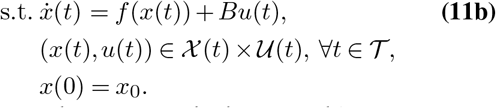

This definition characterizes the best possible moment estimation of the CME according to the given SSA samples and the cost function in Eq. (9) while additionally satisfying the closed moment equations and staying inside the confidence region. Hence, the solution *u*^*^(*t*) is the optimal CC for the moment equations on 𝒯 .

### D. Guaranteed convergent CC using a short-horizon optimization

For an increasing size of the interval 𝒯, the nonlinear optimization problem in Eq. (11) becomes numerically challenging. To overcome this issue, we present in the following a short-horizon optimization based on MPC which is guaranteed to converge to the optimal reachable trajectory (*x*^*^(*t*), *u*^*^(*t*)) without explicitly calculating it. In particular, we use an MPC scheme based on the results in (32), where the authors propose a reference tracking MPC scheme without terminal ingredients for *unreachable* reference trajectories of discrete-time nonlinear systems based on techniques from economic MPC (41, 42). This MPC scheme ensures practical stability of the *unknown* optimal reachable reference (*x*^*^(*t*), *u*^*^(*t*)). Based on this, the results were translated to the case of continuous-time nonlinear systems in (31), which makes it suitable for our setting.

To obtain an optimal estimate of the CME moments which is close to the SSA-based estimation, we want to track the optimal reachable trajectory Eq. (11) without explicitly computing (*x*^*^(*t*), *u*^*^(*t*)). To this end, we define the short-horizon cost function

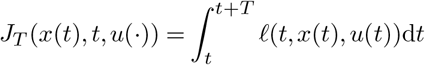

and propose the following: At each time *t* ≥ 0, given the current state *x*(*t*) and an admissible reference 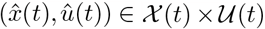, we solve a short-horizon MPC scheme characterized by the optimization problem

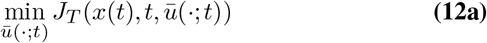

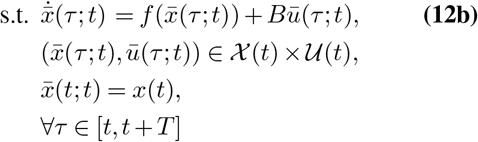

with the prediction horizon *T* satisfying *δ* ≤ *T* ≤ *T*_end_. The optimal trajectory resulting from Eq. (12) is denoted by 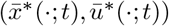, while the control input *u* is chosen as 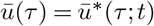 for *τ* ∈ [*t, t* + *δ*). The MPC problem Eq. (12) is solved repeatedly after each sampling period *δ*. The closed-loop dynamics using the MPC controller read then

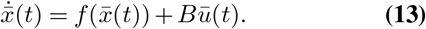

Due to the finite time horizon *T*, the solution to Eq. (12) does in general not exactly follow the optimal trajectory (*x*^*^(*t*), *u*^*^(*t*)). While the application of our proposed CC based on Eq. (12) is generally possible and requires only the definition of some hyperparameters, we can even establish, under mild regularity conditions, recursive feasibility of the MPC problem Eq. (12).

#### Definition 2

The MPC problem Eq. (12) is called *feasible* at time *t* if there exists at least one 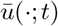 such that the constraints are satisfied. Moreover, the MPC problem Eq. (12) is called *recursively feasible* if initial feasibility at time *t* = 0 implies feasibility of the MPC problem for all *t* ≥ 0.

Further, we show that the resulting closed-loop system practically converges to the optimal reachable reference (*x*^*^(*t*), *u*^*^(*t*)) based on results in (31).To this end, we introduce the notion of practical convergence.

#### Definition 3

A system practically converges to a reference trajectory (*x*^*^(*t*), *u*^*^(*t*)) under control input *μ*(*t*) ∈ 𝒰(*t*) if there exists a *β* ∈ 𝒦ℒ such that the corresponding system state *x* satisfies

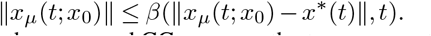

Moreover, the proposed CC recovers the true moments in the limit of infinitely many sample trajectories, which follows from the consistency of the empirical moments estimates. Before stating our main theoretical result, we make the following assumption.

*Assumption 4:* System Eq. (4) representing the MoM is uniformly suboptimally operated off the trajectory *x*^*^, i.e., the following conditions hold (cf. (42, Def. 12)):

- For each *x*_0_ ∈ 𝒳^0^ and each *u* ∈ 𝒰^*∞*^(*x*_0_)

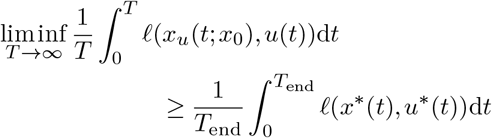

holds, where *U*^*∞*^(*x*_0_) denotes the set of all feasible control sequences of infinite length starting at a given *x*_0_ ∈ 𝒳, and 𝒳^0^ denotes the set of all *x* ∈ 𝒳 such that 𝒰^0^ (*x*) is non-empty.
- There exist 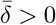 and *d* ∈ 𝒦_*∞*_ such that for each *δ >* 0 and each *ε >* 0 there exists *R*_*ε,δ*_ ∈ ℝ_*≥*0_ such that *δ/R*_*ε,δ*_ ≥ *d (ε)* for all 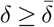 and such that for each *x*_0_ 𝒳^0^ *u*∈ 𝒰 ^*∞*^ (*x*_0_), and each *T* ∈ ℝ_*≥*0_ at least one of the following two conditions hold:

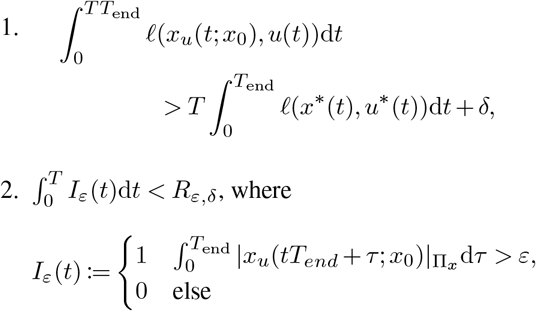

If the minimizer (*x*^*^(*t*), *u*^*^(*t*)) is unique, then Assumption 4 is not restrictive due to the positive definite weight matrix *Q* in the stage cost *𝓁* (compare (32, Dec. IV-D)). The following result establishes the theoretical properties of the proposed CC scheme.

#### Theorem 1

Suppose Assumption 4 holds. Then, there exists a 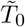 such that for all 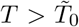the CC 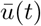 based on the MPC optimization Eq. (12) leads to a moment estimation that practically converges to the *optimal* MoM-based estimates Eq. (11) of the CME and is guaranteed to satisfy physical properties and the given confidence region of the moments. Moreover, the CC recovers the *true* moments for infinitely many sample trajectories, i.e., *N* → ∞.

*Sketch of Proof:* While we omit the proof details here and rigorously prove the theorem in Appendix C, we sketch the main ideas in the following. In particular, the proof relies on practical convergence according to Definition 3, where we use available results from the literature based on Lyapunov stability. To this end, we first exploit that the nonlinear dynamics of the MoM in Eq. (4) are, e.g., polynomial, twice differentiable, and structured such that only the moments of order (*k* + 1) enter (linearly) into the dynamics of the moments of order *k*. Thus, the considered dynamics inherently satisfy necessary assumptions from the literature to guarantee practical convergence. Moreover, the MPC optimization recovers the true moments in the limit *N* since the SSA-based moment estimation converges to the true CME moments for *N* → ∞.

The proposed optimal estimation of the CME moments based on the presented CC scheme is summarized in Algorithm 1. Note that the lower bound 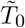 in Theorem 1 is, in general, unavailable since it depends on existent but typically unknown functions. Hence, the theorem shows the existence of a sufficiently long prediction horizon without stating the horizon explicitly. Nevertheless, for a fixed sampling interval *δ*, the closed-loop system stays around the optimal trajectory (*x*^*^(*t*), *u*^*^(*t*)), which is as close as possible to the unreachable reference 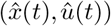, given that a large enough prediction horizon is used. We stress further that when *δ* → 0, *T* should go to infinity (31). Possible future work could use ideas from (43), which presents a nonlinear MPC scheme without terminal ingredients for tracking possibly unreachable setpoints using artificial steady-states (44). In particular, the in (44) proposed results use less restrictive assumptions and might be exploited to indeed derive computable bounds on the necessary prediction horizon, which is beyond the scope of this work.

#### Algorithm 1

Control closure scheme for MoM

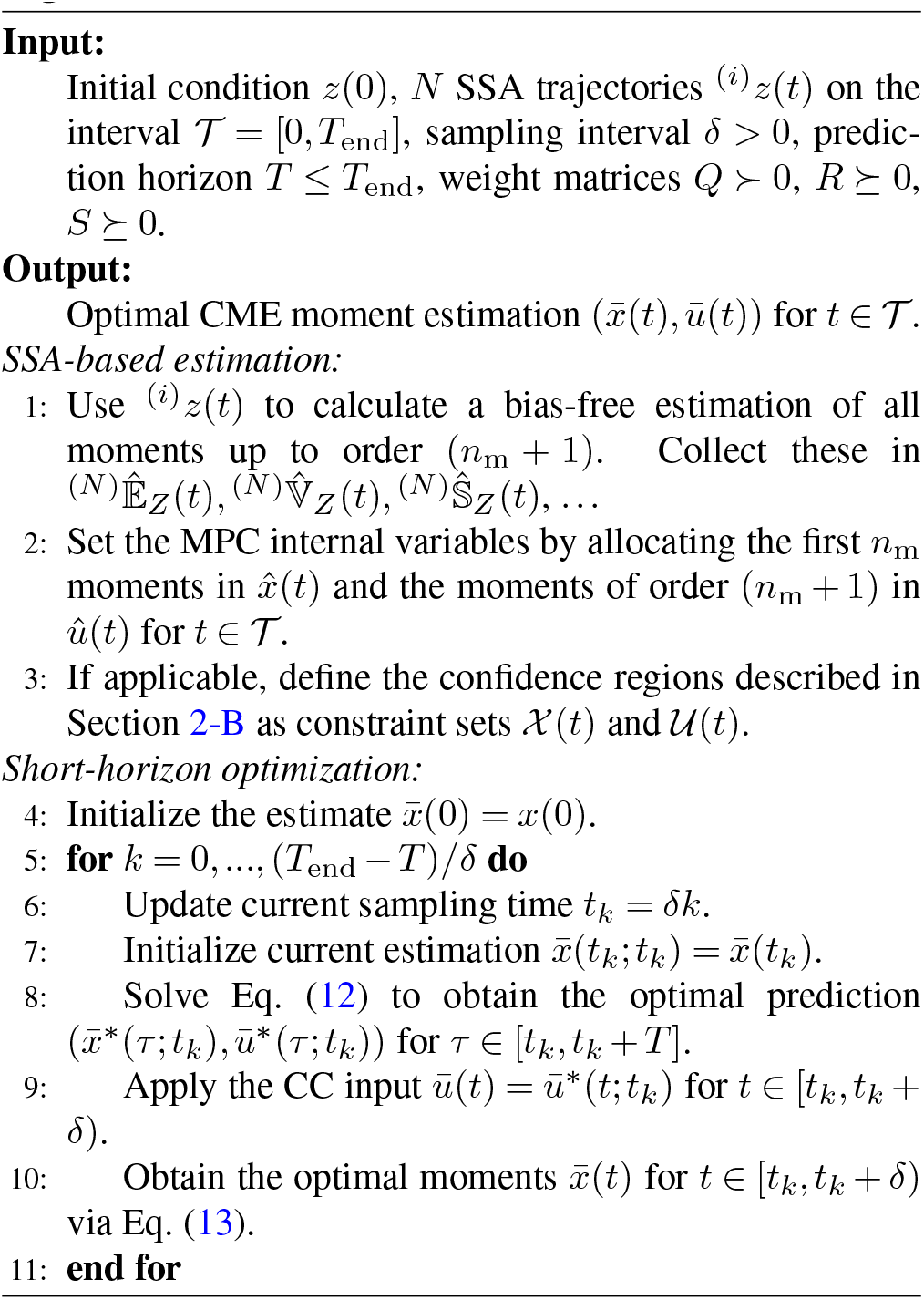

## 3 Simulation examples

In this section, we validate the performance of our CC scheme on three example systems. All considered systems converge to a steady-state distribution on their respective configuration-transition graph, as discussed in (21). Algorithm 1 was implemented in Matlab using MPCTools (45) with its interface to the nonlinear optimization software CasADi (46).

**System 1** The first exemplary system consists of a constant production rate and quadratic degradation and is therefore one of the simplest systems involving quadratic propensities. The reactions read

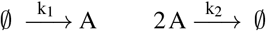

and the MoM ODEs are given by

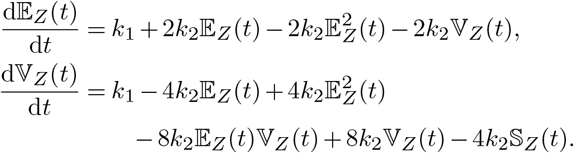

In principle, there is no limit for the number of molecules in System 1 and the space of all possible system configurations Ω_*Z*_ is, therefore, ℕ_0_. In order to still solve the CME numerically, we truncate Ω_*Z*_ to 31 system configurations, leading to Ω_*Z*_ = {0, …, 30}. While this seems to be a very drastic measure, the probability mass not captured by the first 31 configurations is indeed negligible in our setting, which justifies this approximation.

A comparison of the performance of different approaches for moment trajectory estimation for System 1 is shown in Figure 2 for expectation value (top), variance (top second), and skewness (top third). The CME lines represent the true, numeric CME moments. SSA-based moment estimates 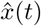 and *û*(*t*) are shown for *N* = 10 and *N* = 100 SSA sample paths. The results of TC fail here completely.

**Fig. 2.**
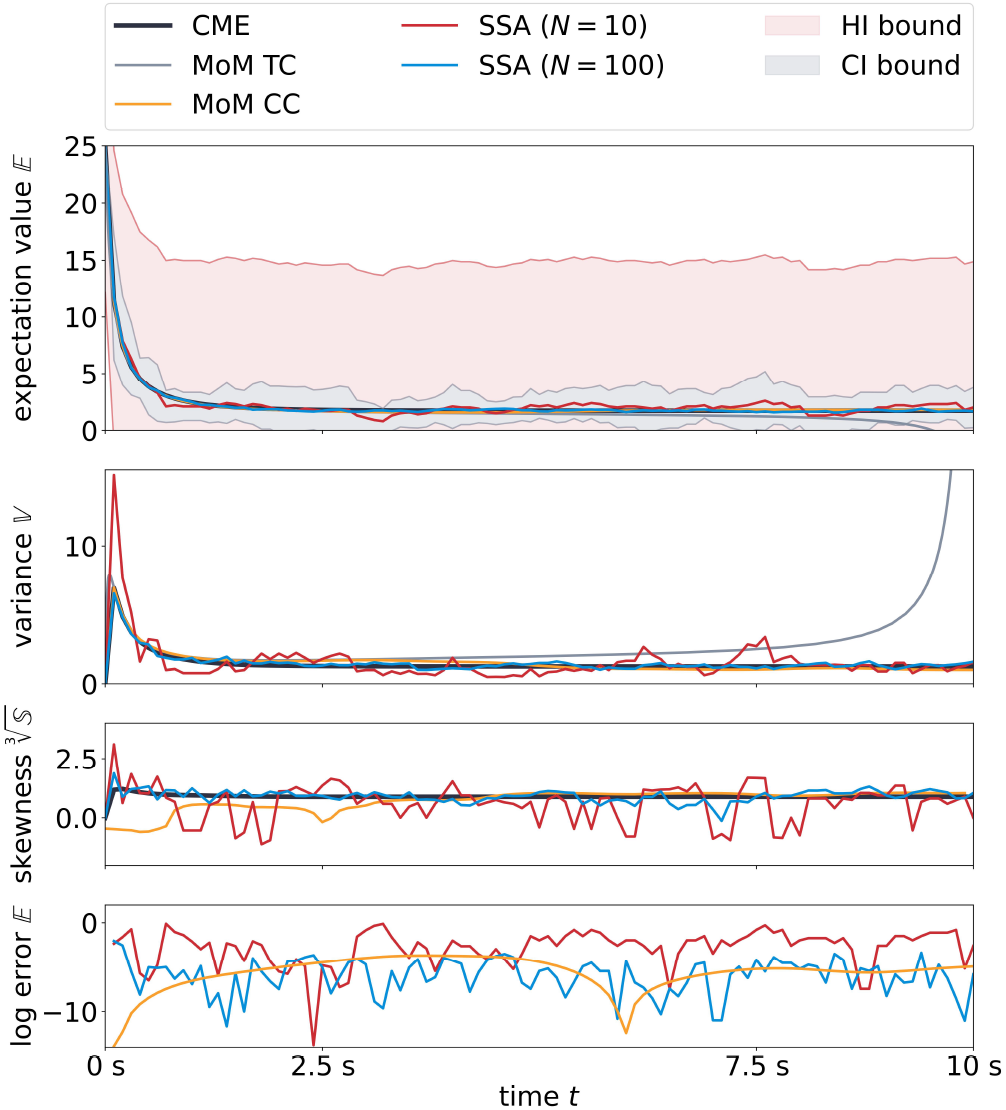
Comparison of moment estimation approaches for System 1. Top and middle. Shown are moment trajectories estimated via SSA samples (*N* = 10 and *N* = 100) and via TC and CC using the average of *N* = 10 SSA trajectories as reference input. The CME reference solution was obtained by numerical integration. Shaded regions are confidence bounds for *N* = 10 obtained with *α* = 0.95. Bottom. Natural logarithm of errors between the true expectation value and its approximations. Results were obtained with initial condition *z*(0) = 25 molecules, which translates to 𝔼_*Z*_ (0) = 25 and 𝕍_*Z*_ (0) = 0 for the moment equations. Reaction rates were set to [*k*_1_, *k*_2_] = [1.25 s^*−*1^, 0.25 s^*−*1^] and the system was simulated over a virtual time of *T*_end_ = 10 s. For CC, we used the parameters *Q* = diag*{*1000, 30*}, R* = 1, *S* = 100, *T* = 1 s, and *δ* = 0.1 s.

The result of our CC scheme utilizes the average over *N* = 10 SSA sample paths as a reference in the objective function.

Confidence regions

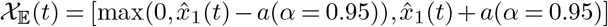

were set for the expectation value by using HI and CI bounds. Please note that the CI bounds require knowledge of the true variance 𝕍_*Z*_ to be valid. As this quantity is typically not available, the CI bounds depicted in Figures 2, 3, and 4 are calculated using the SSA-based estimator 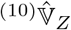 of the true variance and therefore only approximate the true CI bounds. The variance is restricted to non-negative values and the skewness is not constrained in this setting, i.e., 𝒳 (*t*) = 𝒳_𝔼_(*t*) × ℝ_*≥0*_ and 𝒳(*t*) = ℝ. The MPC problem is formulated using the weights *Q* = diag {1000, 300}, *R* = 1, *S* = 100 with the prediction horizon *T* = 1 s and the sampling interval *δ* = 0.1 s. We emphasize that the choice of the weights is not restricted to the values above but also different choices lead to satisfying results. Note that our choice is motivated by the sample accuracy of the SSA estimates, i.e., the variance is roughly weighted by the square root of the weight of the expectation value. Since we do not obtain a reliable estimate for the skewness, the weight *R* is chosen to be small. However, we use the weight *S* = 100*R* to penalize large deviations of the skewness between subsequent time steps to guarantee a certain smoothness. We stress that the absolute values of the weights can be scaled and only the relative relations between *Q, R*, and *S* are important.

**Fig. 3.**
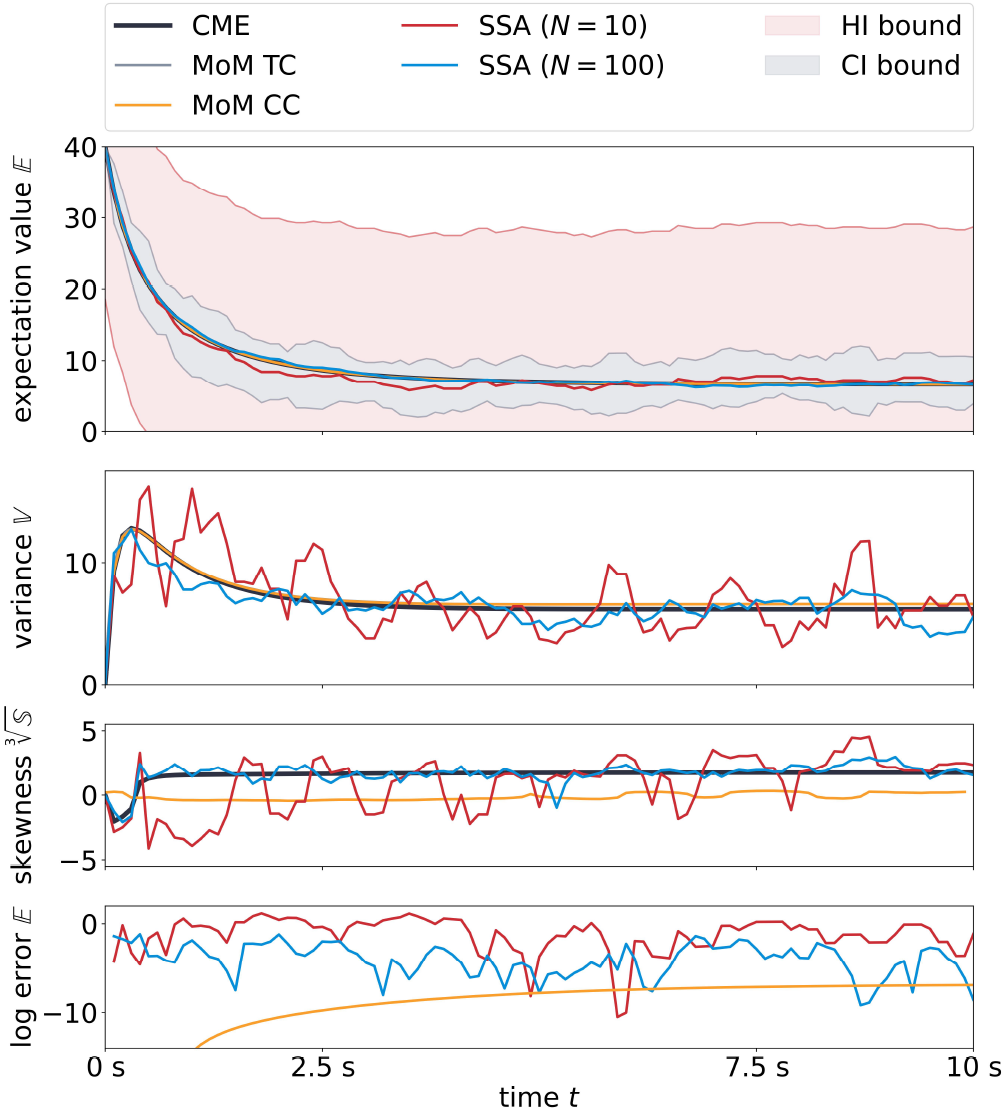
Comparison of moment estimation approaches for System 2. Illustration of results similar to Figure 2. Results were obtained with initial condition *z*(0) = 40 molecules and *c*_A_ = 50 and reaction rates [*k*_1_, *k*_2_] = [0.05 s^*−*1^, 0.025 s^*−*1^]. Similar to Figure 2, we used *Q* = diag {1000, 30 }, *R* = 1, *S* = 100, *T* = 1 s, *δ* = 0.1 s for the CC.

**Fig. 4.**
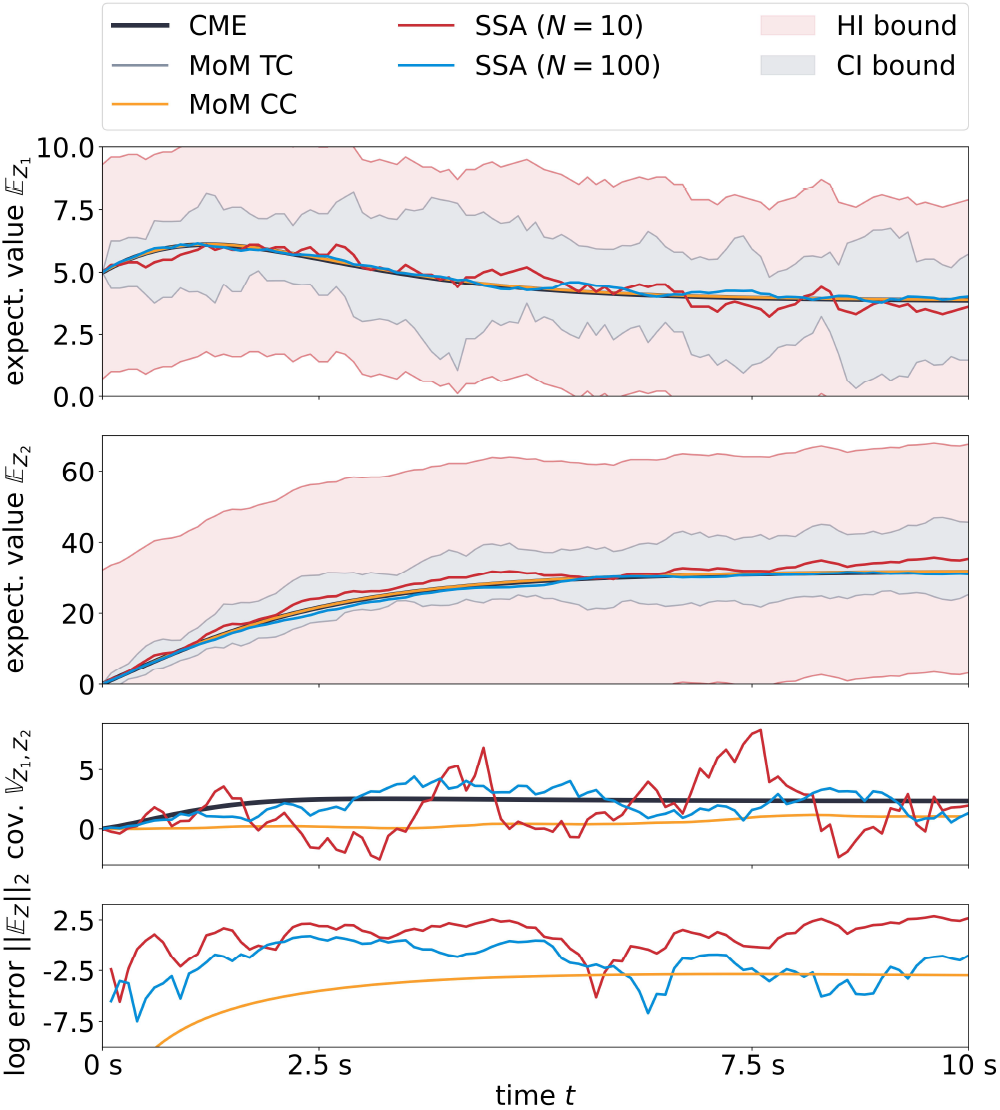
Comparison of moment estimation approaches for System 3. Illustration of results similar to Figure 2. Results were obtained with initial condition *Z*(*t* = 0) = (5, 0) molecules and *c*_A_ = 10 and reaction rates [*k*_1_, *k*_2_, *k*_3_, *k*_4_, *k*_5_, *k*_6_] = [2.0 s^*−*1^, 0.0 s^*−*1^, 0.2 s^*−*1^, 0.2 s^*−*1^, 0.02 s^*−*1^, 0.2 s^*−*1^]. We used *Q* = diag*{*1000, 1000*}, R* = 30, *S* = 100, *δ* = 0.1 s for the CC.

The two first estimated moments resulting from the proposed CC scheme, expectation and variance, are smooth and closely resemble the CME solution. Moreover, the expectation value clearly lies within 𝒳_𝔼_(*t*) for both bounds. The skewness deviates from the reference course for the first 2.5 seconds but approaches the true solution quite well afterward. This becomes especially obvious when considering the squared quadratic error between the true CME solution and its approximations in the bottom sub-graph. Not only is the error of the controlled MoM solution orders of magnitude smaller than the error of the underlying 10-sample SSA approximation, but it is also comparable in magnitude with the SSA approximation for *N* = 100 samples while being significantly smoother at the same time. Further, we note that other closure methods than TC or a higher number of SSA samples might lead to successful estimates of the true CME solutions as well. However, we emphasize that the proposed CC is indeed able to estimate the *true* moments by using only *N* = 10 SSA samples. Additionally, the estimates based on the CC are guaranteed to satisfy the specified SSA-based bounds on the respective moments ensuring proper physical correctness, which is typically not guaranteed for other closure methods.

**System 2** Our second example represents a reversible dimerization reaction

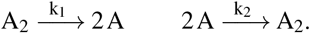

The configuration space of System 2 is limited by a conservation law, 2A_2_(*t*) +A(*t*) = *c*_A_ = 2A_2_(0) +A(0), which can be used to reduce the system to species A only. The MoM ODEs are given by

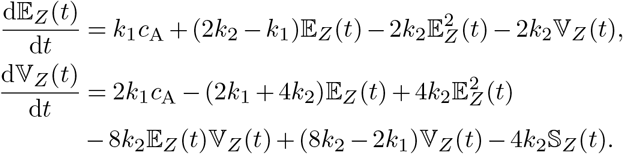

For this simulation example, we chose an initial number of 40 molecules of species A and *c*_A_ = 50, which directly implies the existence of 5 A_2_ molecules for *t* = 0 and Ω_*Z*_ = {0, …, 50}. Moreover, we use the same constraint sets 𝒳(*t*), 𝒰(*t*) and the same optimization parameters *Q, R, S, T, δ* as for System 1. A comparison of results for different approaches is presented in Figure 3 analogously to Figure 2. First and foremost, courses of the true CME moments are extremely well captured by both moment closure approaches, and the expectation value trajectory again lies well within 𝒳_𝔼_(*t*). The reason for the similarity of results of both closure methods is that the control input skewness estimated by the CC scheme is quite close to zero during the whole simulation. In contrast, the SSA-based approximation is as shaky as for System 1.

**System 3** The third exemplary system is chosen to demonstrate the practical applicability of our CC method. To this end, we choose a system that models the self-regulation of the expression of gene A that encodes protein B (47). More specifically, *A*_f_ is the number of free DNA strands, *A*_b_ is the number of DNA strands with a bound protein and B is the number of free proteins. The governing reactions read

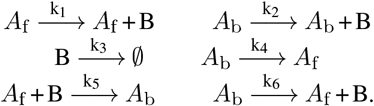

Similar to System 2, the three molecule species can be encoded by a two-dimensional state *Z* = (A_f_, B) with the help of a conservation law, as the sum of free and bound DNA is constant, i.e., A_f_(*t*) + A_b_(*t*) = *c*_A_ = A_f_(0) + A_b_(0). It is practically unrealistic to close the MoM using potentially all four skewness terms. Thus, the MoM in this application example predicts the expectation values of both chemical species and uses the covariance 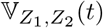 as the closure moment (*n*_m_ = 1), but we note that the CC is not restricted to this simplification. Indeed, due to the nature of the involved reactions, the variances 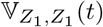 and 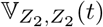 of both individual species do not have an impact on the respective expectation values, as they do not arise when deriving the MoM equations

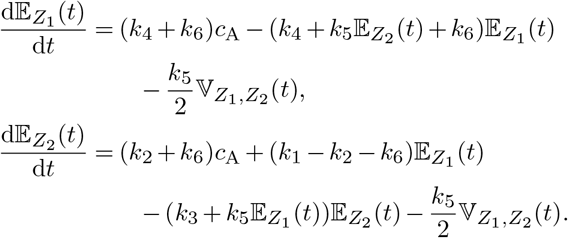

The number of proteins B is generally unbounded. Similar to our treatment of System 1, we truncate 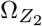 to 76 system configurations leading to 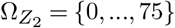. Again, this truncation does not neglect any probability mass beyond numeric tolerances. The results depicted in Figure 4 are created with initial numbers of *Z*(*t* = 0) = (5, 0) molecules and *c*_A_ = 10. As we did for Systems 1 and 2, we used sample-based reference trajectories for the expectation value and the variance. Please keep in mind, that in contrast to the previous systems, this means that we also employ a reference trajectory for the control input. Moreover, we use the confidence regions

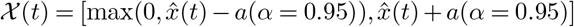

for both expectation values, similar to 𝒳_𝔼_ for System 1 and System 2, and 𝒰(*t*) = ℝ for the covariance. The parameters for the MPC optimization are chosen as *Q* = diag {1000, 1000}, *R* = 30, *S* = 100 with prediction horizon *T* = 1 s and sampling interval *δ* = 0.1 s. We note that the weight matrices are strongly related to the ones used for System 1 and System 2. Again, the weight *R* = 30 of the covariance is roughly the square root of the weights *Q*_11_ = *Q*_22_ = 1000 of the expectation values.

Similar to the previous figures, Figure 4 visualizes the performance of different approaches for moment trajectory estimation for System 3. In contrast to the previous figures, however, we now depict two expectation values in the top plots, each with its own confidence regions 𝒳(*t*). The third sub-graph again depicts the time course of the closing moment, which in this case is the covariance. Lastly, the bottom sub-graph shows the natural logarithm of the 2-norm of the error of the expected numbers of molecules 𝔼_*Z*_(*t*). The results arising from this gene self-regulatory system align with previous results: TC and the proposed CC scheme provide smooth solutions of the MoM that are in addition very accurately resembling the true CME moments. As guaranteed by construction, the CC solution stays within its specified bounds and converges to the true CME solution as *T* → ∞.

## 4 Conclusion

Applying the MoM to approximate the CME’s moments is a challenging task due to the interdependence structure of the considered moments. In this work, we interpreted the system of MoM ODEs in the framework of control theory to present a non-heuristic closure scheme that provably stays close to the true solution in a probabilistic sense. More specifically, our CC scheme draws a few SSA samples from the true CME solution and uses the corresponding sample-based moment estimates as a physically meaningful reference trajectory. The closure moments are then used as the system’s control input in order to follow the SSA trajectories while still obeying the MoM dynamics. Indeed, the CC leads to physically meaningful results that closely approximate the true moments even when TC fails to do so. Moreover, it outperforms purely sample-based approximation techniques as it smooths the sample averages according to the MoM dynamics. The MoM and hence basically all moment closure approaches require the considered moments of the CME solution to exist for the time interval of interest, which we have assumed (and numerically tested) in this work. However, it should be noted that this is a delicate issue since reaction systems with solutions whose moments are not defined can easily be constructed (48). These systems can show wildly stochastic behavior, and in particular, SSA-based moment estimates do not converge for increasing sample sizes. Moment-based methods terribly fail when applied to such systems without caution. Generally, this restriction has to be kept in mind when dealing with the MoM and related approaches. We demonstrated the capabilities of CC using two representative test-bed systems and a gene self-regulatory reaction network. The latter emphasizes that our closure scheme can in principle be applied to systems with several molecular species. In addition, it is straightforward to consider higher-order moments, as indicated by the more general dimensions at several places in the manuscript. However, a severe limitation is that the number of moments scales very badly with the number of molecular species in the system. For *n*_c_ chemical species, we have *n*_c_ expectation values, an *n*_c_ × *n*_c_ covariance matrix, 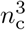 skewness values, and the number grows likewise for higher moments. This is true for other moment approaches as well, but also affects the computation time in our case via repeated optimization. Finally, we note again that a particular strength of our method compared to other existing closure schemes is that we are able to provide probabilistic guarantees that the calculated moments stay in a neighborhood of the true moments and hence cannot arbitrarily diverge. Moreover, our scheme provides several tuning points to control this distance. First, the number of SSA samples correlates negatively with the size of the confidence intervals. Second, different weights can be given to different moments, and the smoothness of the input can be regulated via hyperparameters in the MPC objective function. Finally, computational effort can to some extent be regulated by appropriately choosing the horizon in the short-horizon MPC scheme. Overall, we believe that CC has great potential to be widely applied to describe the moments of biochemical reaction systems.

## Funding

Funded by Deutsche Forschungsgemeinschaft (DFG, German Research Foundation) under Germany’s Excellence Strategy - EXC 2075 – 390740016 and within grant AL 316/15-1 – 468094890. We acknowledge the support by the Stuttgart Center for Simulation Science (SimTech). V. Wagner and R. Strässer thank the Graduate Academy of the SC SimTech for its support.

## Abbreviations

CC: Control Closure
CI: Chebyshev-Inequality
CLT: Central Limit Theorem
CME: Chemical Master Equation
HI: Hoeffding’s Inequality
MoM: Method of Moments
MPC: Model Predictive Control
ODE: Ordinary Differential Equation
SSA: Stochastic Simulation Algorithm
TC: Truncation Closure

## Availability of data and materials

All code is available via https://fairdomhub.org/models/846.

## Competing interests

The authors declare that they have no competing interests.

## Authors’ contributions

N.R. and F.A. supervised the project and acquired funding, V.W. and R.S. implemented the examples, all authors contributed to the theoretical results, R.S., V.W., and N.R. wrote the manuscript, all authors reviewed the manuscript.

## Acknowledgments

This preprint was formatted using a LATEX class by Ricardo Henriques that can be accessed here.

## A Method of Moment ODEs

For Systems 1 and 2, we are interested in the expectation value 𝔼_*Z*_(*t*) and variance 𝕍_*Z*_(*t*) of the time-dependent number of system molecules *Z*(*t*). To derive the MoM equations, we start with Proposition 2.5 in (3). For the special case of one chemical species, the equations read

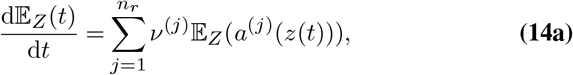

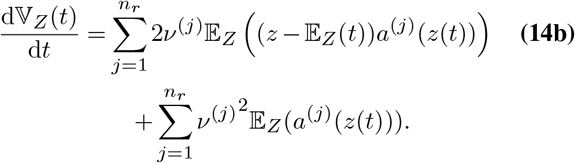

A crucial next step is a reasonable approximation of 𝔼_*Z*_(*a*^(*j*)^(*z*(*t*))). To this end, we assume that the system’s propensities are perfectly approximated by a second-order Taylor approximation. This assumption is fulfilled for all mass-action kinetics, as their propensities are polynomials of order two or less. More specifically, our assumptions imply

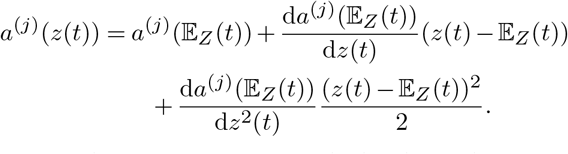

Applying the expectation value to both sides of the equation results in the desired characterization

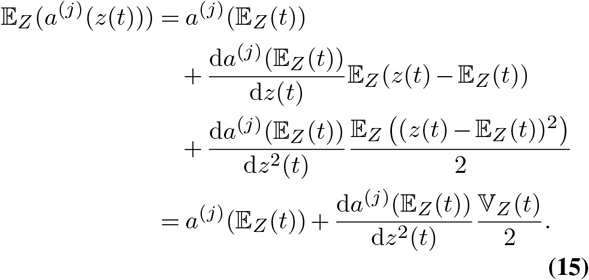

By inserting Eq. (15) into the original MoM ODEs Eq. (14), we obtain the final MoM equations

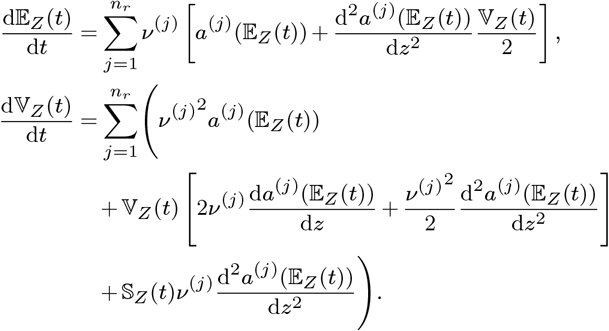

## B Deriving a probabilistic upper bound for SSA-based moment estimation from the CLT

The theorem of Berry-Esseen (34, 35) is an improved version of the CLT. We use it to derive a probabilistic bound for the expectation estimation error of a sample average.

### Theorem 2

(Berry-Esseen) For *a* ∈ R and *N* samples, consider the cumulative density *F*_*N*_ of the shifted and scaled sample average 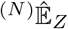

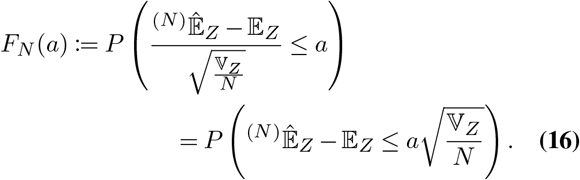

Then, we conclude

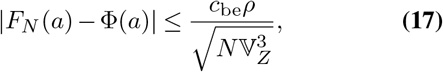

where Φ is the standard-normal cumulative distribution.

We derive the probabilistic bound by first reformulating the quantity of interest and using the definition of *F*_*N*_ . Subsequently, Theorem 2 is employed to achieve an upper bound. After this, terms are rearranged and we introduce the error function that is related to Φ by the equality

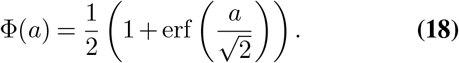

Altogether, the derivation reads as

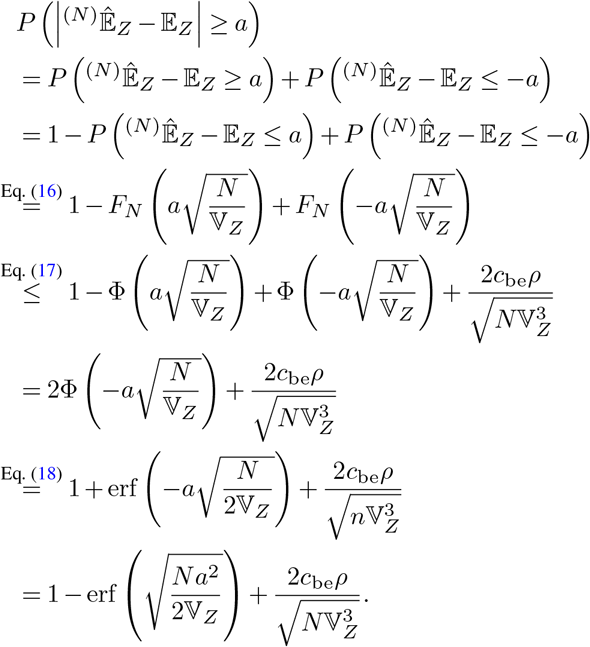

## C Theoretical properties

In this section, we provide the theoretical background on MPC which is necessary to establish the theoretical guarantees of our CC scheme in Theorem 1. More precisely, we present theoretical guarantees for the class of polynomial systems Eq. (4), including all mass-action systems, leading to probability bounds for the error between true and estimated moment trajectories. In particular, we present results on closed-loop (practical) stability of the closed-loop Eq. (13) based on (31). To this end, we recall the notion of practical stability.

### Lemma 3

((49, Thm. 2.4), (30, Thm. 2.23)) Let *V* be a time-varying practical Lyapunov function with

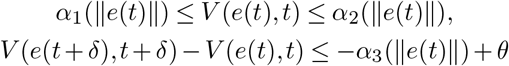

for all *V* (*e*(*t*), *t*) ≤ *V*_max_, 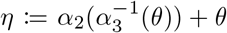 and *α*_1_, *α*_2_, *α*_3_ ∈, *𝒦θ >* 0. Then, for all initial conditions *V* (*e*(0), 0) ≤*V*_*max*_, the origin *e* = 0 is uniformly practically asymptotically stable, i.e., the system uniformly converges to the set {(*e, t*) | *V* (*e, t*) ≤ *η}* as *t →* ∞.

In the following, we establish practical stability of system Eq. (4) and show that the MPC problem in Eq. (12) is recursively feasible. To this end, we need the following local incremental stabilizability assumption (31, 32).

*Assumption 5:* There exist a continuous control law *κ* : 𝒳× 𝒳× 𝒳 → ℝ^*m*^, a *δ*-Lyapunov function *V*_*δ*_ : 𝒳× 𝒳 × 𝒰 → ℝ_≥0_, that is continuous in the first argument and satisfies *V*_*δ*_(*ξ, ξ, v*) = 0 for all (*ξ, v*) ∈ 𝒳× 𝒰, and positive parameters *c*_*δ,l*_, *c*_*δ,u*_, *δ*_loc_, *k*_max_, *ρ*, such that for all initial conditions (*x*(0), *ξ*(0), *v*(0)) ∈ 𝒳 × 𝒳 × 𝒰 with *V*_*δ*_(*x*(0), *ξ*(0), *v*(0)) ≤ *δ*_loc_ hold

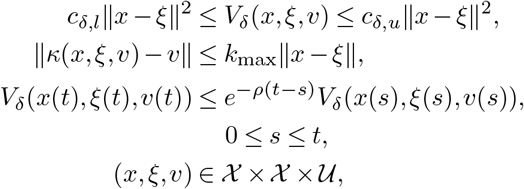

with

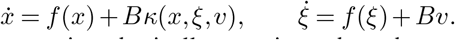

This assumption basically requires that the controller *κ*(*x, ξ, v*) leads to perfect tracking of (*ξ, v*) when *x* is sufficiently close to *ξ*. The constant *k*_max_ serves as a local Lipschitz bound on the nonlinear control law *κ*.

As our goal is to stay close to the reference trajectory 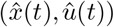, the MPC scheme should track the *unknown* reference (*x*^*^(*t*), *u*^*^(*t*)). To this end, the subsequent analysis assumes compactness of 𝒳× 𝒰, which is satisfied for bounded state space Ω_*Z*_. Since Subsection 2-B only discusses probabilistic bounds of the expectation, but no bounds for higher-order moments, this requirement is generally violated for unconstrained moments. However, some system classes allow also the derivation of probabilistic bounds for higher-order moments and, alternatively, it is always possible to define a large enough interval around the SSA-based moment estimate which then contains the true moments with high probability and leads to the compactness of 𝒳× 𝒰. Hence, using additionally the boundedness of the SSA-based estimate 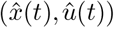 for the considered systems, the difference between the reference and the optimal reachable trajectory is bounded as well. We make the following assumption to guarantee constraint satisfaction.

*Assumption 6:* The optimal reachable reference trajectory (*x*^*^(*t*), *u*^*^(*t*)) from Definition 1 is such that *V*_*δ*_(*x*(*t*), *x*^*^(*t*), *u*^*^(*t*)) ≤ *δ*_r_ implies

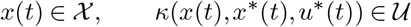

with *V*_*δ*_, *κ* from Assumption 5 and some *δ*_r_ *>* 0. There exists a function *α*_*V*_ ∈ 𝒦 such that *V*_*δ*_(*x*(*t*), *x*^*^(*t*), *u*^*^(*t*)) ≤ *δ*_r_ and *V*_*δ*_(*y, x*^*^(*t*), *u*^*^(*t*)) ≤ *δ*_r_ imply

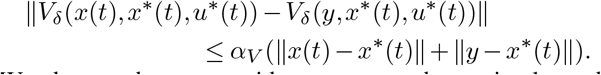

We denote the error with respect to the optimal reachable reference by *e*_r_(*t*) = *x*(*t*) − *x*^*^(*t*) and the stage cost of the optimal reachable reference trajectory by *𝓁*^*^(*t*) = *𝓁*(*t, x*^*^(*t*), *u*^*^(*t*)). Further, we make the following dissipativity assumption (31, 32, 50).

*Assumption 7:* There exists a time-varying storage function *λ* : ℝ× → ℝ such that for all (*x*(*t*), *u*(*t*)) ∈ 𝒳× 𝒰 the rotated stage cost *L* satisfies

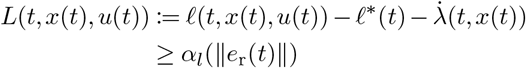

for all *t* ≥ 0, *δ >* 0, with *α*_*l*_ ∈ 𝒦_*∞*_. Furthermore, for all *x*(*t*) ∈ 𝒳 the storage function is uniformly bounded by

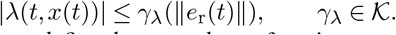

Moreover, we define the rotated cost function

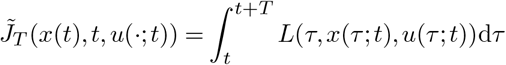

and the corresponding rotated MPC problem

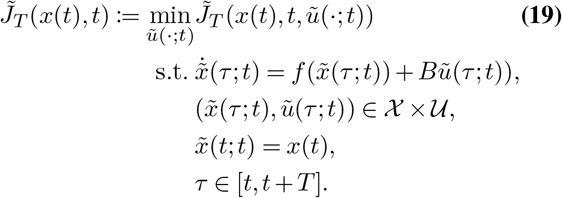

We make the following assumption to establish a lower bound on the optimal cost.

*Assumption 8:* For any given positive constant *δ* ≤ *T*, there exists a function *α*_*δ*_ ∈ 𝒦 satisfying that

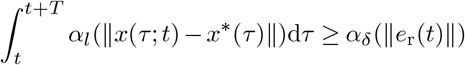

for all feasible *x*(*τ* ; *t*) ∈ 𝒳, *τ* ∈ [*t, t* + *T*], where 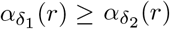 if *δ*_1_ *δ*_2_.

For practical tracking, we introduce the set

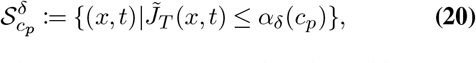

with *α*_*δ*_ according to Assumption 8 and *c*_*p*_ defined by

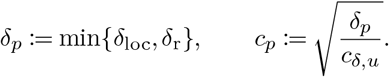

### Theorem 4

((31, Thm. 3.1)) Let Assumptions 5, 6, 7, and 8 be satisfied. Then, there exists a 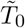 and a function 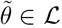, such that for all 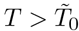 and all initial conditions satisfying 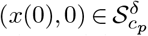, the MPC problem Eq. (12) is recursively feasible and the closed-loop system Eq. (13) satisfies

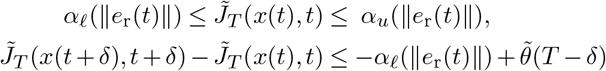

for *t* ≥ 0. The set 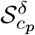 is positively invariant and the origin *e*_r_ = 0 is uniformly practically asymptotically stable under the closed-loop dynamics Eq. (13).

*Proof:* This result is taken from (31) and we omit the proof for conciseness.

The resulting closed-loop system practically tracks the *unknown* optimal reachable reference (*x*^*^(*t*), *u*^*^(*t*)). We note that the assumption on the initial condition in Theorem 4 can be relaxed by requiring a longer time-horizon 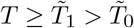 (see (31, Thm. 3.2)).

*Remark 9:* Note that the results in (31, 32) are only valid for *periodic* references. Even though in our case the reference is *non*periodic, we can still use the presented results to ensure closed-loop stability over any finite interval 𝒯 (compare (32, Rem. 8)). More precisely, any reference 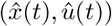 on the interval 𝒯 can be extended to a periodic reference 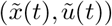, where 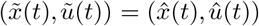 for all *t* ∈ 𝒯 and the period length *p ≥T*_end_. Then, clearly, both reference trajectories with the same initial condition cannot be distinguished on the finite interval 𝒯.

Now we can prove Theorem 1 establishing theoretical guarantees for our proposed CC scheme estimating the moments of the CME.

*Proof of Theorem 1:* We rely on the notion of practical stability according to Definition 3, which is ensured by Lemma 3, and the result of Theorem 4. To this end, we first need to show that the considered setting satisfies all assumptions of Theorem 4. Note that the nonlinear dynamics in Eq. (4) are twice differentiable. Further, the MoM dynamics Eq. (4) are such that the linearized system (*A*(*ξ*), *B*) with 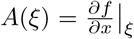 is locally controllable around any point (*ξ, v*) ∈ 𝒳 × 𝒰. This results from the particular structure of the dynamics, where the moments of order (*k* + 1) enter only linearly in the dynamics of the moments of order *k*, i.e., *A*_*j,j*+1_(*ξ*) = *A*_*j,j*+1_ ∈ ℝ\{0} for all *j* = 1, …, *n* − 1 and 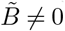. Thus, for *n*_c_ = 1 and *n*_m_ = 2 we have

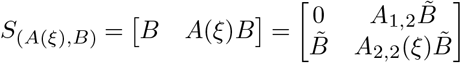

with *S*_(*A*(*ξ*),*B*)_ having full rank for all *ξ*, i.e., the linearized system is controllable. Hence, Assumption 5 is satisfied for system Eq. (4) (cf. (31, Rem. 2.1), (32, Prop. 1)). As discussed in (41, 51), Assumption 6 requires local continuity of the *δ*-Lyapunov function *V*_*δ*_ around the optimal reachable reference trajectory (*x*^*^(*t*), *u*^*^(*t*)) which is also ensured by local controllability of the system. Computing suitable storage functions for Assumption 7 is typically challenging for general nonlinear systems with arbitrary reference trajectories 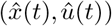. However, the existence of such a storage function can be shown assuming local controllability (see (52, Def. 3.6.4)) and that the system is uniformly suboptimally operated off the trajectory *x*^*^ according to Assumption 4 (compare (32, Lm. 4)). Thus, Assumption 7 is satisfied for system Eq. (4) with Assumption 4. Moreover, as noted in (31), a lower bound satisfying Assumption 8 can be constructed for polynomial systems and, thus, in particular also for system Eq. (4), using convex optimization (53). Hence, all the assumptions of Theorem 4 are satisfied.

It remains to show that the initial condition satisfies 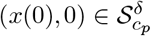 with 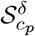 defined in Eq. (20). Since we assume in our setting exact knowledge of the initial condition, i.e., 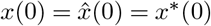 and *e*_r_(0) = *x*(0) − *x*^*^(0) = 0, we obtain 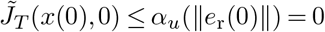 due to (31, Prop. 3.1), where 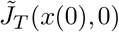 denotes the optimal rotated cost function defined in Eq. (19). Hence, the initial condition (*x*(0), 0) of our setting is always contained in the set 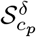, and, thus, we can directly apply Theorem 4 to show the first result of Theorem 1 for the constraint set 𝒳× 𝒰 consisting of the SSA-based confidence bounds combined with physical properties of the considered moments.

The second claim results from the fact that the SSA-based moment estimation 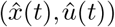 converges to the true CME moments for *N* → ∞. Since the MoM is exact if we know the true closure moments and, in the limit, the considered reference 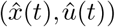 matches with the true moments and is thereby compatible with the dynamics Eq. (4), the MPC optimization also follows the true CME moments for *N* → ∞.

## Notes

### Competing Interest Statement

The authors have declared no competing interest.

### Summary of Updates

Added a third example system with 2 species in Section 3 System 3. More coherent notation throughout the paper.

